# Autophagy promotes programmed cell death and corpse clearance in specific cell types of the *Arabidopsis* root cap

**DOI:** 10.1101/2022.02.16.480680

**Authors:** Qiangnan Feng, Riet De Rycke, Yasin Dagdas, Moritz K. Nowack

## Abstract

Autophagy is a conserved quality control pathway that mediates the degradation of unnecessary or dysfunctional cellular components by targeting them to the lysosomes or central vacuoles. Autophagy has been implicated in the regulation or execution of regulated cell death processes in a wide range of eukaryotes. However, its function in developmentally controlled programmed cell death (dPCD) in plants remains little studied and controversial. Here, we investigated the role of autophagy in dPCD using the Arabidopsis root cap as an accessible and genetically tractable model system. We show that autophagic flux is induced prior to dPCD execution in both root cap tissues, the columella and the lateral root cap (LRC), and impaired in autophagy-deficient mutants. These mutants show a strongly delayed cell death and an absence of corpse clearance in the columella during and after their shedding into the rhizosphere. However, autophagy deficiency does not affect dPCD execution or corpse clearance in LRC cells at the distal end of the root cap. Our results demonstrate that autophagy promotes dPCD in a highly cell-type specific manner, and present the root cap as a powerful model system to study organ-specific autophagy *in vivo*.

## Introduction

Regulated cell death (RCD) describes genetically encoded mechanisms to dispose of cells in a tightly controlled way. Different RCD processes are vital for growth and development, immunity and stress responses across eukaryotes. In animals, a growing number of RCD subroutines have been described^1^ that can act in two opposing contexts: On the one hand, RCD occurs as integral part of development, tissue turnover, and ageing, largely independent of environmental perturbation. These physiological forms of RCD are often referred to as programmed cell death (PCD). On the other hand, RCD can occur as response to infection and abiotic stresses^1^.

In plants, these two contextual categories have been described as developmental PCD (dPCD) and environmental PCD (ePCD)^2^. One of the most studied ePCD types is the hypersensitive response (HR), a localized plant host response at the site of pathogen infection^3^. In the context of plant development, dPCD processes occur as the last step of differentiation in particular cell types, at the end of organ senescence, or facultatively as the result of cellular signalling^4^. Despite the multiple and often indispensable functions of dPCD for plant growth and reproduction, still comparatively little is known about its molecular regulation.

Over the last years, the *Arabidopsis thaliana* (Arabidopsis) root cap has been established as a model system to investigate dPCD in its native developmental context *in planta*. The root cap surrounds the root meristem stem cell niche, and is important to sense environmental cues, including gravity, for directed root growth^5^. In Arabidopsis, dPCD is an integral part of root cap differentiation, restricting root cap organ size to the root meristem^6^. The root cap consists of two distinct tissues originating from different stem cells: the proximally positioned columella that is located at the very root tip, and the lateral root cap (LRC) that flanks the root meristem up to its distal end at the start of the root elongation zone^5^. Incipient continuous layers of columella and LRC cells are generated by repeated stem cell division close to the quiescent centre, displacing the older root cap layers to the root periphery. The root cap organ size is kept constant by a combination of dPCD and shedding of cells into the rhizosphere^7,8^: When epidermal cells start to elongate at the distal end of the meristem, their neighbour cells of the distal LRC (Figure 1A) undergo a tightly controlled dPCD and corpse clearance process on the root surface, limiting the root cap extent to the meristematic zone^6^. Eventually, cell death advances proximally towards the columella creating distinct PCD sites that demark the respective end of each LRC layer^6^. Finally, the columella cells, together with the adjacent proximal LRC cells, get shed as cell packages into the rhizosphere^9^, where they undergo dPCD shortly thereafter^10^.

**Figure 1.**
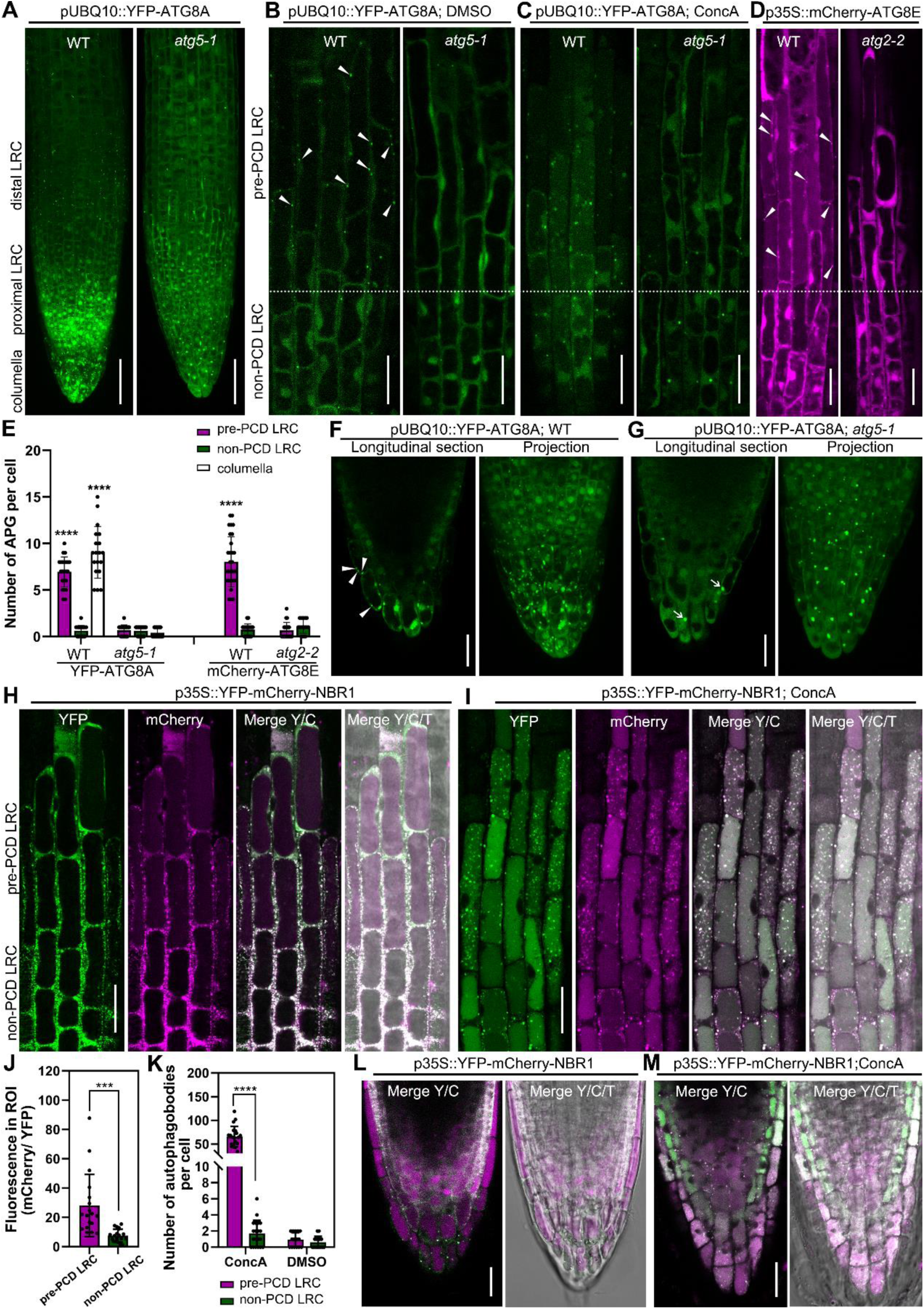
Visualization of the dynamics of autophagosomes/autophagy-related structures in root cap cells. (**A-C**), Confocal laser scanning micrograph (CLSM) of LRC cells from seedlings at 4 days after germination (DAG) expressing pUBQ10::YFP-ATG8A in the wild type and the *atg5-1* mutant, projection of the root tip (A), longitudinal section of LRC cells treated with DMSO as control (B), or treated with 1 μM ConcA for 8 h (C). Scale Scale bars are 50 μm (A), 20 μm (B-C). (**D**), CLSM of lateral root cap (LRC) cells from 4 DAG seedlings of expressing p35S::mCherry-ATG8E in wild type and *atg2-2* mutant. Scale bars are 20 μm. (**E**), Quantification of autophagosomes in LRC and columella cells. Results are means ± SD (> 20 counted cells from several separate seedlings). **** indicates a significant difference (*t* test, P < 0.0001). (**F-G**), CLSM of columella cells from 4 DAG seedlings expressing pUBQ10::YFP-ATG8A in the wild type (E) and the *atg5-1* mutant (F). Scale bars are 20 μm. White arrow heads indicate autophagosomes, white arrows indicate YFP aggregates. (**H-I**), CLSM of LRC cells from 4 DAG seedlings expressing p35S::YFP-mCherry-NBR1 in the wild type (H), treated with 1 μM ConcA for 8 h (I). Scale bars are 20 μm. (**J**), Quantification of the vacuolar import of NBR1 in LRC cells as shown in (H). Quantification was performed by dividing the average gray value in cytosol (YFP) with the average gray value in the vacuole (mCherry), using split channel images in ImageJ. The graphical representation is based on about 20 counted cells from several separate seedlings, with standard deviation indicated as Scale bars. *** indicates a significant difference (*t* test, P < 0.001). (**K**), Quantification of autophagic bodies in LRC cells treated with 1 μM ConcA for 8 h as shown in (I). The graphical representation is based on 20 cells from several separate seedlings, with standard deviation indicated as Scale bars. **** indicates a significant difference (*t* test, P < 0.0001). (**L-M**), CLSM of columella cells from 4 DAG seedlings expressing p35S::YFP-mCherry-NBR1 in wild type (L), treated with 1 μM ConcA for 8 h (M). Scale bars are 20 μm. Y/C indicates merge of the YFP channel and the mCherry channel; Y/C/T indicates merge of the YFP channel, the mCherry channel, and the transmission channel.

Developmental PCD has been characterized as a succession of cellular processes including PCD preparation, execution, and corpse clearance^4^. The NAC (no apical meristem [NAM], Arabidopsis thaliana activation factor [ATAF], cup-shaped cotyledon [CUC]) family transcription factors SOMBRERO/ANAC033 (SMB), ANAC046 and ANAC087 have been shown to orchestrate the preparation for dPCD execution and corpse clearance in the root cap. These transcription factors promote the expression of dPCD-associated genes^6,10,11^. While *smb* mutants show an extended life span of distal LRC cells in the root elongation zone, *anac046* mutants exhibit an extended life span of the columella and the adjacent proximal LRC cells during and after their shedding into the rhizosphere^6,10^. Despite these studies unravelling the role of transcriptional networks in root cap dPCD, the cellular pathways that execute cell death remain poorly understood.

In animals, types of RCD that rely on components of the autophagic machinery have been described as autophagy-dependent RCD^1^. Autophagy is a bulk degradation process that delivers unnecessary or dysfunctional cellular components to the lysosome for degradation. Autophagic responses often occur as stress-adaptation response, hence acting generally in rather a cytoprotective than a cytotoxic way. However, in specific developmental and pathophysiological settings, the autophagic machinery can contribute to cellular demise^12^.

A major autophagy pathway, macroautophagy, (referred to as “autophagy” hereafter) is largely conserved in plants^13^. Cytoplasmic cargo is engulfed by so-called phagophores that close to form double-membrane bound autophagosomes. Proteins encoded by the conserved autophagy-related (ATG) genes ATG9, ATG2 and ATG18 mediate phagophore formation. Later, the phagophore gets decorated with ATG8, a process involving ATG5 and ATG7^13^. The autophagosome transports its cargo to central vacuole, where the outer membrane of the autophagosome fuses with the vacuolar membrane (tonoplast), while the remaining single-membrane structure, the autophagic body, is taken up by the vacuole. Finally, autophagic bodies and their cargo are broken down by vacuolar hydrolases^13-15^. Arabidopsis mutants such as *atg7-1*^16^, *atg5-1*^17^, and *atg2-2*^18^ are deficient in autophagosome formation and autophagic flux to the vacuole. These mutants are hypersensitive to nutrient-limiting conditions, suggesting that autophagy plays a key role in nutrient recycling as adaptation to such conditions. In addition, several *atg* mutants exhibit an early leaf senescence phenotype^16,17^.

In analogy to the situation in animals, the involvement of autophagy in plant PCD has been controversially discussed^19^. While some studies reported a pro-survival role of autophagy, for instance in leaf senescence, immunity, or lace plant leaf development^20-22^, others suggested a pro-death role, e.g. in xylem formation^23^, suspensor elimination^24^, or plant immunity^25^. These data suggest that the function of autophagy is complex and can play either “pro-death” or “pro-survival” roles depending on the experimental settings and biological situation. Therefore, understanding the function of autophagy especially in the context of dPCD will benefit from further study.

Here we used the root cap model to investigate the relevance of autophagy in dPCD. We show that autophagic flux increases prior to dPCD in both LRC and columella cells, and that autophagic flux is inhibited in *atg* mutants. Intriguingly, the life span of *atg* mutant columella cells in the rhizosphere is significantly extended, and post-mortem corpse clearance does not occur. By contrast, the lack of autophagy does not affect dPCD execution or corpse clearance in the distal LRC. We show that root cap autophagy phenotypes can be produced in a root cap-autonomous manner, and do not depend on the established NAC transcription factor pathways. Our results demonstrate a dPCD-promoting function of autophagy in the columella, but not in the distal LRC, suggesting that autophagy roles in dPCD can considerably diverge between different cell types of the same plant organ.

## Results

### Autophagic flux is increased prior to dPCD in root cap cells

To test the involvement of autophagy in the root cap dPCD process, we investigated autophagosome/autophagic body formation and autophagic flux in wild-type and *atg* mutant root cap cells. To this end, we imaged root cap cells of Arabidopsis lines carrying pUBQ10::YFP-ATG8A or p35S::mCherry-ATG8E constructs^26,27^. In the wild-type, we detected YFP-ATG8A signal in both the cytosol as well as in autophagic foci in LRC cells preparing for PCD (“pre-PCD” LRC cells). By contrast, we found predominantly cytosolic YFP signal in LRC cells that are not yet preparing for PCD (“non-PCD cells”, Figure 1A, B). In the autophagy-defective *atg5-1* mutant^17^, we counted significantly less YFP-ATG8A foci in “pre-PCD” cells (Figure 1E), indicating that autophagy is activated prior to LRC cell death in an ATG5-dependent manner. These results were confirmed in wild-type seedlings expressing an alternative autophagy marker, p35S::mCherry-ATG8E (Figure 1D, E), suggesting our findings represent a general autophagy response.

The Arabidopsis root cap organ consists of two different tissues, the LRC and the columella^8^. To analyze the autophagy related structures in columella cells, we imaged plants expressing pUBQ10::YFP-ATG8A. We found YFP signal within small punctate foci of columella cells in the outermost root cap layer in wild type roots (Figure 1F). By contrast, YFP-ATG8A signal was free cytosolic and clustered in large cytoplasmic aggregates in the columella cells of *atg5-1* mutants, as previously described (Figure 1G)^27^. Quantification demonstrated that the number of small punctate foci (autophagosomes) in mature columella cells is significantly increased compared to “non-PCD” cells (Figure 1E). This suggests that similar to the “pre-PCD” LRC cells preparing for cell death, autophagic activity is also increased in mature columella cells. Concanamycin A (ConcA) is an established drug to neutralize vacuolar pH by inhibiting vacuolar-type ATPases^28^. Elevated vacuolar pH results in stabilized autophagic bodies, allowing to quantify autophagic delivery to the vacuole. After 8 hours of ConcA treatment, numerous YFP-ATG8A-positive autophagic bodies accumulated in wild type “pre-PCD” LRC cells (Figure 1C), and mature columella cells (Movie S1). Conversely, we detected no vacuolar autophagic bodies in ConcA-treated *atg5-1* mutant, consistent with autophagy deficiency in this mutant (Figure 1C and Movie S2). To monitor and quantify autophagic flux to the vacuole, we imaged root cap cells expressing a double-tagged autophagic cargo, YFP-mCherry-NBR1. This reporter is delivered to the vacuole by autophagy where low pH allows to differentiate between cytoplasmic to vacuolar fluorescence ratios comparing the pH-sensitive YFP and the less pH-sensitive mCherry ^29^. We identified numerous NBR1 foci in the cytosol, and an increasingly strong vacuolar mCherry fluorescence in “pre-PCD” LRC and columella cells (Figure 1H, J and L). This suggests that autophagic flux is increased in root cap cells prior to PCD execution. To confirm this result, we treated the double-tagged NBR1 line with ConcA. In the central vacuole of both the LRC and the columella, there are more autophagic bodies in “pre-PCD” root cap cells than in “non-PCD” root cap cells (Figure 1I, K and M). These results clearly demonstrate that autophagy activation is associated with root cap dPCD and that autophagic flux is increased in root cap cells prior to PCD execution.

### Autophagy is required for timely dPCD onset of proximal LRC and columella cells

To investigate the roles of autophagy in root cap PCD, we performed a live-death assay using fluorescein diacetate (FDA) and propidium iodide (PI) staining of roots from 5-day old wild type and *atg* mutant seedlings.

The fluorescence of FDA indicates cell viability, and PM permeabilization was visualized by PI entry into the cell which designated the time point of cellular death^10^. We found the longitudinal root cap extent (“root cap size”) and distal LRC PCD onset in *atg* mutants to be indistinguishable from the wild type (Figure 2A and 2B). LRC PCD is followed by rapid cell corpse clearance on the root surface^6^. Used time-lapse imaging to monitor the PI-positive LRC cell corpse clearance we found no differences between wild type and *atg2-2* mutant cells (Figure S1). This result is consistent with the absence of PI-stained cellular remnants of distal LRC cells in the elongation zone of *atg* mutants (Figure 2A) that can be found in mutants delayed in LRC post-mortem corpse clearance^6,10^. However, while the proximal LRC and some columella cells in the wild type had already undergone PCD prior to shedding, the same cells in *atg* mutants remained alive as indicated by FDA fluorescence and absence of PI staining (Figure 2A). Detailed analysis of optical sections confirmed that significantly more proximal LRC and columella cells were alive in the *atg* mutants (Figure 2C and D). To investigate if root cap development was delayed in *atg* mutants, we performed an FDA and FM4-64 double staining to visualize both viability and cell outlines in root tips of 5-day old seedlings. Root cap development and architecture was indistinguishable between wild type and *atg* mutants, each had five well-organized root cap layers (Figure S2A). Moreover, statolith-formation as a hallmark of columella differentiation^30,31^ was indistinguishable in the wild type and *atg* mutants (Figure S2B-F). Interestingly, statoliths in 5-day old *atg* mutant seedlings appears smaller and more numerous than in the wild type. However, statolith degradation prior to cell death occurred independently of autophagy and showed no difference in *atg* mutants and the wild type. These results indicate that autophagy is not critical for root cap development and statolith degradation, but that autophagy promotes timely dPCD initiation in proximal LRC and columella cells.

**Figure 2.**
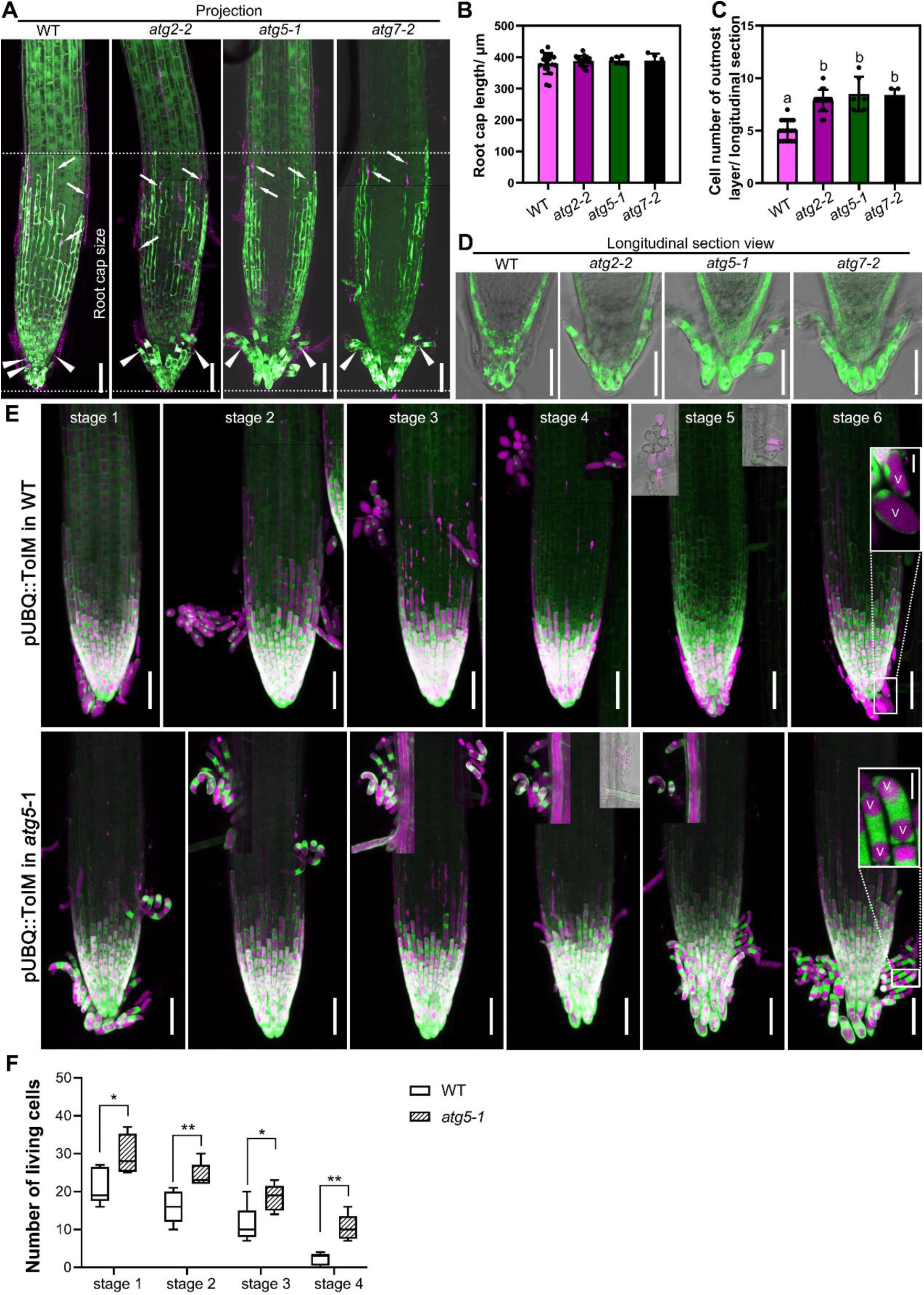
Autophagy is required for dPCD onset of proximal LRC and columella cells. (**A**), CLSM (z-stack) of root tips from 5 DAG seedlings of wild type, *atg2-2, atg5-1* and *atg7-2*, pulse labeled with FDA and PI. White arrows indicate PI-stained nuclei in LRC cells. Scale bars are 50 μm. (**B**), Quantification of root cap size of wild type and *atg* mutants. Results are means ± SD. (**C**), Cell number quantification of the outmost root cap layer of wild type and *atg* mutants on longitudinal sections. Results are means ± SD. Each mutant is significantly different from the wild type, as indicated by different letters (one-way ANOVA, Dunnett’s multiple comparison test, P < 0.05). (**D**), Longitudinal section of wild type and *atg* mutants, pulse labeled with FDA. Single optical section in (D) and maximal z section projection in (A) are generated from the same root. Scale bars are 50 μm. (**E**), pUBQ10::ToIM expression in the wild type and *atg5-1* was analyzed every 12 h (stages 1-6), with the visual onset of root cap shedding defined as stage 1. *atg5-1* mutant shows longer living shed basal LRC and columella cells compared with the wild type. EGFP signal is shown in green in the cytoplasm, and mRFP signal is shown in magenta in the vacuole. Scale bars are 50 μm. Close-up of cells are inserted in stage 6. V indicates the vacuole. Scale bars are 10 μm for inserted images. (**F**), Quantification of living, shed cells from wild type and *atg5-1* mutant at four stages after shedding shows that columella cells of the mutant lived longer compared with the wild type. Box plot whiskers show minimum and maximum values; * indicates a significant difference (*t* test, P < 0.05). ** indicates a significant difference (*t* test, P < 0.01).

To be able to analyze the shedding root cap cells in a time course analysis, we introduced a tonoplast integrity marker (ToIM) controlled by the pUBQ10 promoter^10^ into the *atg5-1* mutant. The ToIM consists of a cytoplasmic eGFP marker and a vacuolar mRFP marker, which makes it possible to visualize vacuolar collapse as a hallmark of PCD when both signals merge^6^. By analyzing ToIM-expressing wild type and *atg5-1* mutant growing in imaging chambers at 12 h intervals for several days, we did not observe any aberration in the shedding process of *atg5-1* root cap cells. However, we observed that *atg5-1* root cap cells remained viable significantly longer than the wild-type ones after shedding into the rhizosphere (Figures 2E and F). Already in stage 1 (defined by the visual onset of columella cell separation in 5-day old seedlings), more living root cap cells were found in the mutant compared with the wild type, indicating that in the wild type, PCD of some columella cells is already executed before shedding. Also, in stage 2 and 3 (12 h and 24 h after stage 1, respectively), significantly more root cap cells were viable in the *atg5-1* mutant than in the wild type. Even in stage 4 and 5 (36 h and 48 h after shedding), living root cap cells could still be detected in the *atg5-1* rhizosphere, which was rarely observed in wild-type plants. Shedding of the next-younger root cap layer started in stage 6 (60 h after stage 1), again showing substantially more living proximal LRC and columella cells in the *atg5-1* mutant in comparison to the wild type (Figure 2E). Interestingly, the vacuolar morphology was altered in the long-lived *atg5-1* root cap cells; instead of a single large vacuole observed in the wild type, most root cap cells showed two smaller vacuoles at either end of the cell (Figure 2E).

To examine the ultrastructure of these long-lived root cap cells, we investigated *atg2-2* mutant by transmission electron microscopy (TEM). In both the wild type and *atg2-2* mutant, the plasma membrane, a dense cytoplasmic matrix, numerous mitochondria, and well-organized Golgi stacks were clearly discernible (Figure S3D-F and J-L). TEM confirmed the altered vacuole structure in *atg* mutants (Figure S4J and K).

In sum, our results demonstrate that autophagy plays a clear PCD-promoting role in the context of dPCD execution in shedding and shed proximal LRC and columella cells.

### Autophagy is crucial for cell-autonomous corpse clearance in columella cells

To investigate a putative involvement of autophagy in the cell-autonomous corpse clearance that occurs during and after dPCD, we cultivated *atg* mutants and wild type seedlings for 14 days on vertical agar plates. This method of cultivation provides minimal friction between the growing root tip and the growth medium, allowing for the accumulation of several dead root cap cell layers. Using FDA/PI staining, we found and increased number of living FDA-positive root cap cells in *atg* mutants, confirming our results on younger seedlings (Figure 3A, Figure S3A). Additionally, we counted significantly more PI-stained columella cell corpses in *atg* mutants compared with the wild type (Figure 3A and B). This indicates that PI-stained cellular corpses are efficiently degraded in the wild type, while cell corpse clearance is impaired in *atg* mutants. Time-lapse imaging revealed that corpse clearance in *atg5-1* mutant columella cells is completely inhibited, or at least strongly delayed: In wild type, PI-stained nuclei were cleared within 24 h, while in *atg5-1* mutant, PI positive nuclear remnants appeared practically unchanged as late as 60 h after PI entry (Figure 3C and D). TEM imaging confirmed these results: In the wild type, cellular remnants in dead cells appeared strongly condensed and organelles were in various stages of degradation (Figure S3G, H and I). By contrast, in the *atg2-2* mutant, columella cell corpses did not collapse, were lined by a continuous plasma membrane, and contained abundant remains of cellular contents (Figure S3M, N and O).

**Figure 3.**
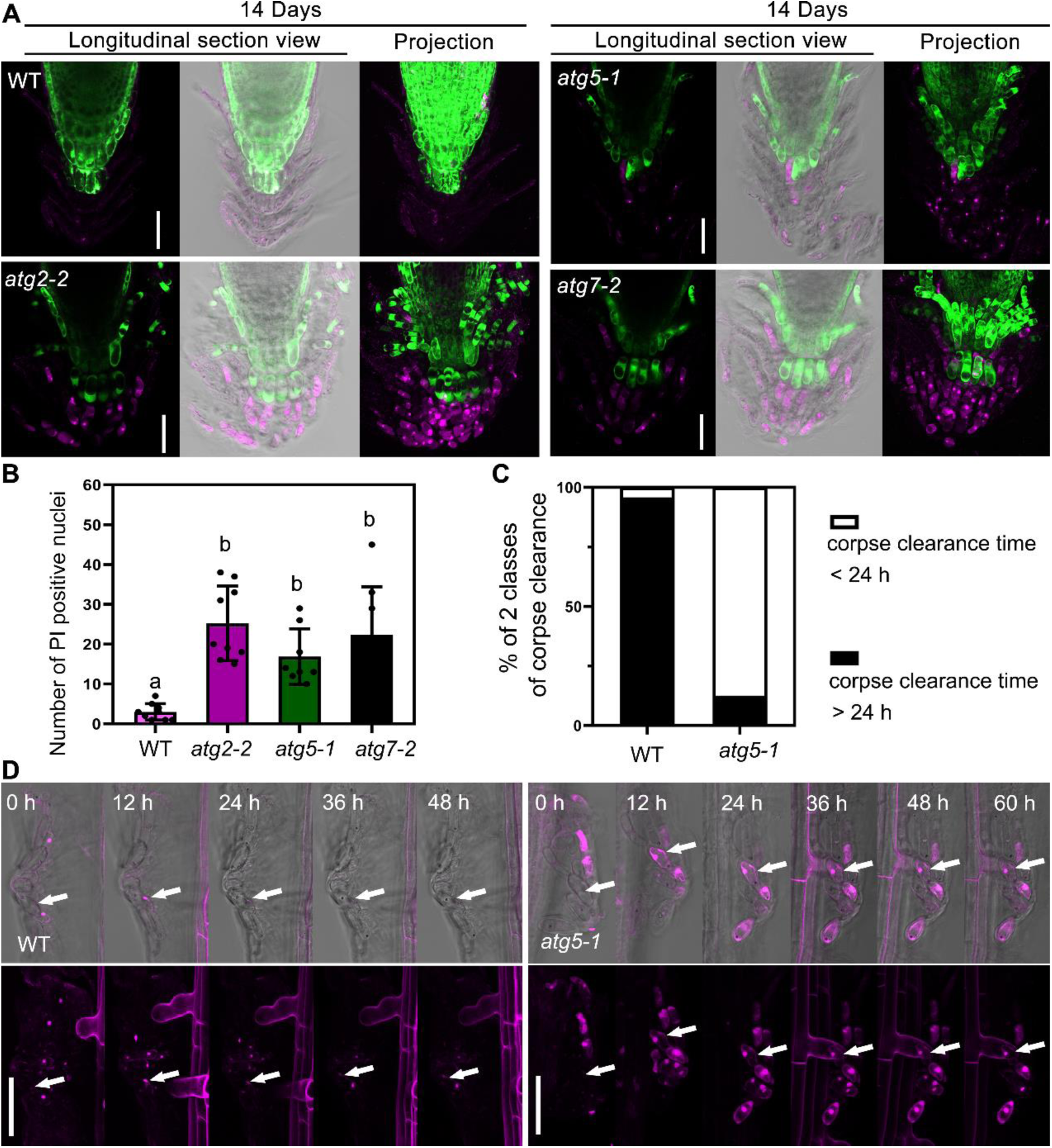
Autophagy promotes the postmortem nuclear degradation in columella cells. (**A**), CLSM of root tips from 14 DAG seedlings of wild type, *atg2-2, atg5-1* and *atg7-2*, pulse labeled with FDA and PI. Scale bars are 50 μm. (**B**), Quantification of PI-stained nuclei of wild type and *atg* mutants, shown in (A). Results are means ± SD. Each mutant is significantly different from the wild type, as indicated by different letters (one-way ANOVA, Dunnett’s multiple comparison test, P < 0.05). (**C**), Quantification of corpse clearance of columella cells. Nuclei of *atg5-1* took longer to be degraded after cell death compared with nuclei from the wild type. Box plots show data from at least 40 nuclei from at least 6 different roots. (**D**), Kymograph showing the delay of nuclear degradation in the *atg5-1* mutant compared with the wild type. Cells were imaged every 12 h. PI is shown in magenta, and cells are shown with (up row) and without (down row) transmitted light. The PI signal is shown as a z-stack projection, whereas the bright-field channel is shown as a single stack. Scale bars are 20 μm.

### Root cap-specific loss of function and complementation confirms tissue-inherent functions of autophagy

As autophagic processes are active throughout tissues of the entire plant, we wondered whether the changes observed are due to loss of autophagy in the root cap specifically, or possibly a secondary consequence of an organism-wide absence of autophagic processes. To differentiate between these possibilities, we first used a clustered regularly interspaced short palindromic repeats (CRISPR)-based tissue-specific knockout system, CRISPR-TSKO, that expresses a fluorescently tagged Cas9 exclusively in the root cap^32^. Combined with guide-RNAs (gRNAs) targeting key ATG genes we intended to generate transgenic lines in which only the root cap is defective in autophagy. To verify the efficiency of our constructs, a pSMB::Cas9-GFP expression cassette was combined with two gRNAs targeting ATG2 or ATG5 (pSMB::Cas9-GFP;ATG2 or pSMB::Cas9-GFP;ATG5), and transformed into a homozygous Arabidopsis line expressing the autophagy reporter p35S::mCherry-ATG8E. In “pre-PCD” LRC cells, both constructs produced a significant reduction in the number of mCherry-labelled cytoplasmic foci and a clear reduction of vacuolar mCherry signal (Figure 4A and B), indicating a defect in autophagosome formation and delivery of ATG8E to the vacuole. The same constructs were transformed into the wild type for phenotype analysis, and four independent transgenic lines of each construct were selected and analyzed. All lines showed significantly delayed corpse clearance in columella cells of 14-day old seedlings as indicated by PI staining, comparable to the conventional *atg* mutants (Figure 4C-4F). Additionally, FDA staining revealed a pronounced columella- and proximal LRC-cell longevity phenotype, similar to the one of the *atg2-2* mutant (Figure S4).

**Figure 4.**
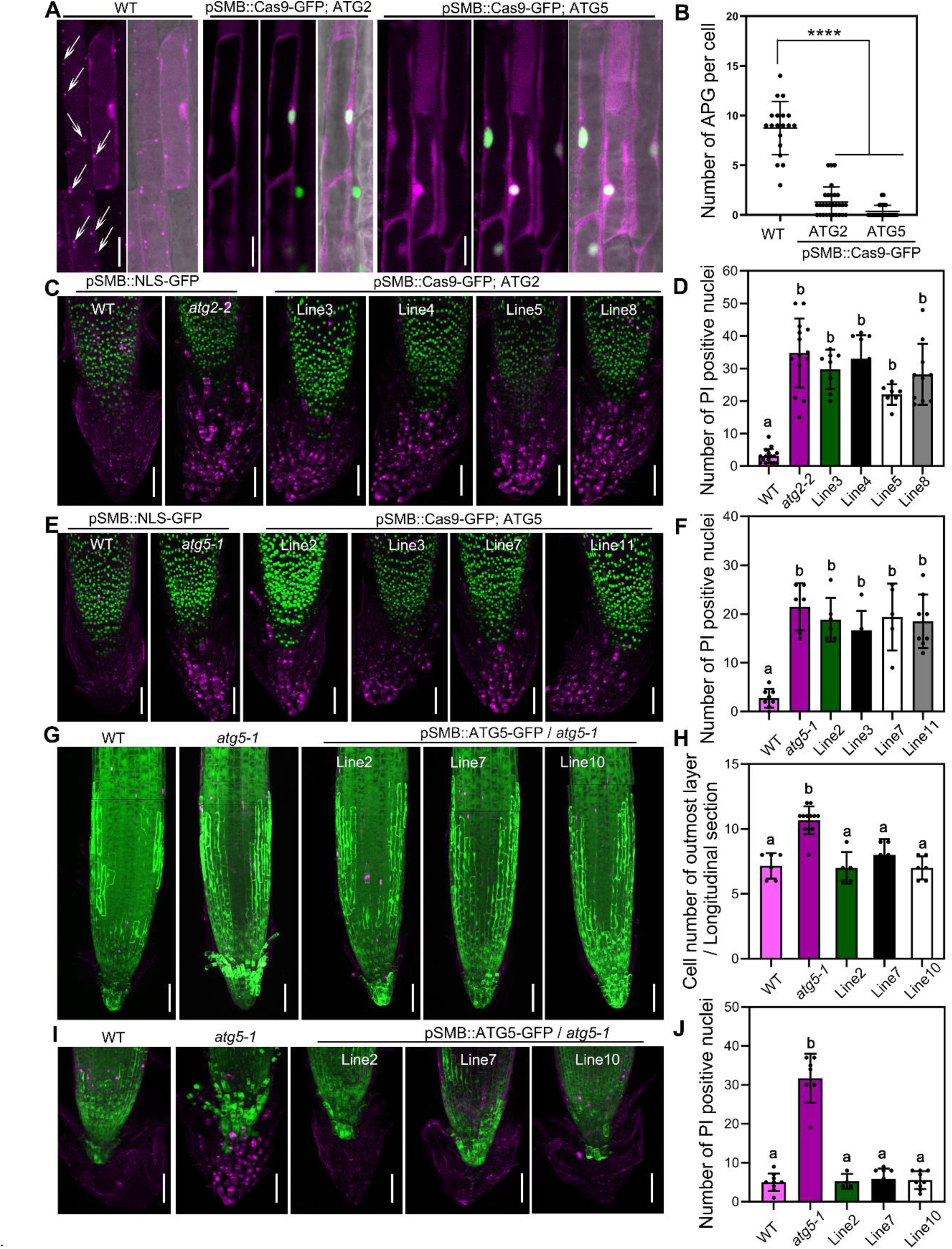
Autophagy controls cell death in a root-cap autonomous fashion. (**A**), CLSM of LRC cells from 4 DAG seedlings expressing p35S::mCherry-ATG8E in wild type, and in the CRISPR-TSKO lines pSMB::Cas9-GFP;ATG2 and pSMB::Cas9-GFP;ATG5. mCherry signal is shown in magenta, GFP signal is shown in green. White arrows indicate autophagosomes. Scale bars are 20 μm. (**B**), Quantification of autophagosome in LRC cells. Results are means ± SD (> 20 counted cells from several separate seedlings). **** indicates a significant difference (Student’s *t* test, P < 0.0001). (**C**), CLSM (z-stack projection) of root tips from 14 DAG seedlings of pSMB::NLS-GFP in wild type, in *atg2-2*, and four different lines of pSMB::Cas9-GFP;ATG2, pulse labeled with PI (magenta). Scale bars are 50 μm. (**D**), Quantification of PI-stained nuclei of seedlings shown in (C). Results are means ± SD. Each mutant is significantly different from the wild type, as indicated by different letters (one-way ANOVA, Dunnett’s multiple comparison test, P < 0.05). (**E**), CLSM (z-stack projection) of root tips from 14 DAG seedlings of pSMB::NLS-GFP in wild type, in *atg5-1*, and four different lines of pSMB::Cas9-GFP;ATG5, pulse labeled with PI (magenta). Scale bars are 50 μm. (**F**), Quantification of PI-stained nuclei of seedlings shown in (E). Results are means ± SD. Each mutant is significantly different from the wild type, as indicated by different letters (one-way ANOVA, Dunnett’s multiple comparison test, P < 0.05). (**G**), CLSM (z-stack) of root tips from 5 DAG seedlings of wild type, *atg5-1* and three different root-cap specific complementation lines (pSMB::GFP-ATG5) in the *atg5-1* mutant, pulse labeled with FDA and PI. Scale bars are 50 μm. (**H**), Cell number quantification of the outmost root cap layer shown in (G) on longitudinal sections. Results are means ± SD. Each mutant is significantly different from the wild type, as indicated by different letters (one-way ANOVA, Dunnett’s multiple comparison test, P < 0.05). (**I**), CLSM (z-stack projection) of root tips from 14 DAG seedlings of wild type, in *atg5-1*, and three different root-cap specific complementation lines (pSMB::GFP-ATG5) in the *atg5-1* mutant, pulse labeled with PI (magenta). Scale bars are 50 μm. (**J**), Quantification of PI-stained nuclei of seedlings shown in (I). Results are means ± SD. Each mutant is significantly different from the wild type, as indicated by different letters (one-way ANOVA, Dunnett’s multiple comparison test, P < 0.05).

Finally, we generated transgenic lines expressing a root-cap specific ATG5 complementation construct under the pSMB promoter in the *atg5-1* mutant background. Three independent lines showed a complete rescue of the longevity and corpse clearance phenotypes of the *atg5-1* mutant (Figure 4G-J). These results demonstrate an autonomous regulation and function of autophagy in the root cap context.

### Autophagy occurs independent of established dPCD gene regulatory networks

Loss of key autophagy regulators caused a columella longevity phenotype reminiscent of the double mutant *ana046 anac087*, two NAC transcription factors that jointly control the timely onset of PCD in the proximal root cap^10^. Hence, we investigated whether autophagy is controlled by, or controlling, these transcription factors. To this end, we subjected 5- or 14-day old *anac046 anac087* mutant seedlings to FDA/PI staining. As previously reported^10^, the double mutant exhibits delayed corpse clearance in distal LRC cells (Figure S5F). In the proximal root cap, we confirmed a significantly increased number of living cells comparable to the one found in *atg* mutants (Figure S5F and G). Also, we detected a delay in corpse clearance in the *anac046 anac087* mutant in the form of a significant increase in PI-stained nuclear remnants compared with the wild type (Figure S5H and I), which had not been reported previously. However, we identified clear differences between *anac046 anac087* and *atg* mutants: While the corpse clearance in autophagy mutants is strongly delayed or even completely absent (Figure 3), in *anac046 anac087* mutants the degradation is merely delayed, leading to a lower number of accumulated PI-positive cells (Figure S5H and I). In addition, the vacuolar morphology of *anac046 anac087* mutants is similar to wild type in proximal LRC cells, and not reminiscent of the one of *atg* mutants (Figure S5J).

To investigate if the upregulation of autophagic activity prior to root cap cell death is regulated by ANAC046 and ANAC087, we transformed a GFP-ATG8E autophagy reporter into *anac046 ana087* mutants, along with the Col-0 wild type and *atg5-1* mutants as controls. In line with our earlier results on YFP-ATG8A and mCherry-ATG8E (Figure 1), GFP-ATG8E shows an increase of autophagic activity and autophagic flux to the vacuole in root cap cells preparing for dPCD, while this activity is comprised in *atg5-1* mutants (Figure S5A). However, the increased autophagic activity in *anac046 anac087* mutant root cap cells preparing for dPCD is indistinguishable from the one of the wild type (Figure 5A-C) These results indicate that the activation of autophagy in aged root cap cells occurs independently of ANA046 and ANAC087.

**Figure 5.**
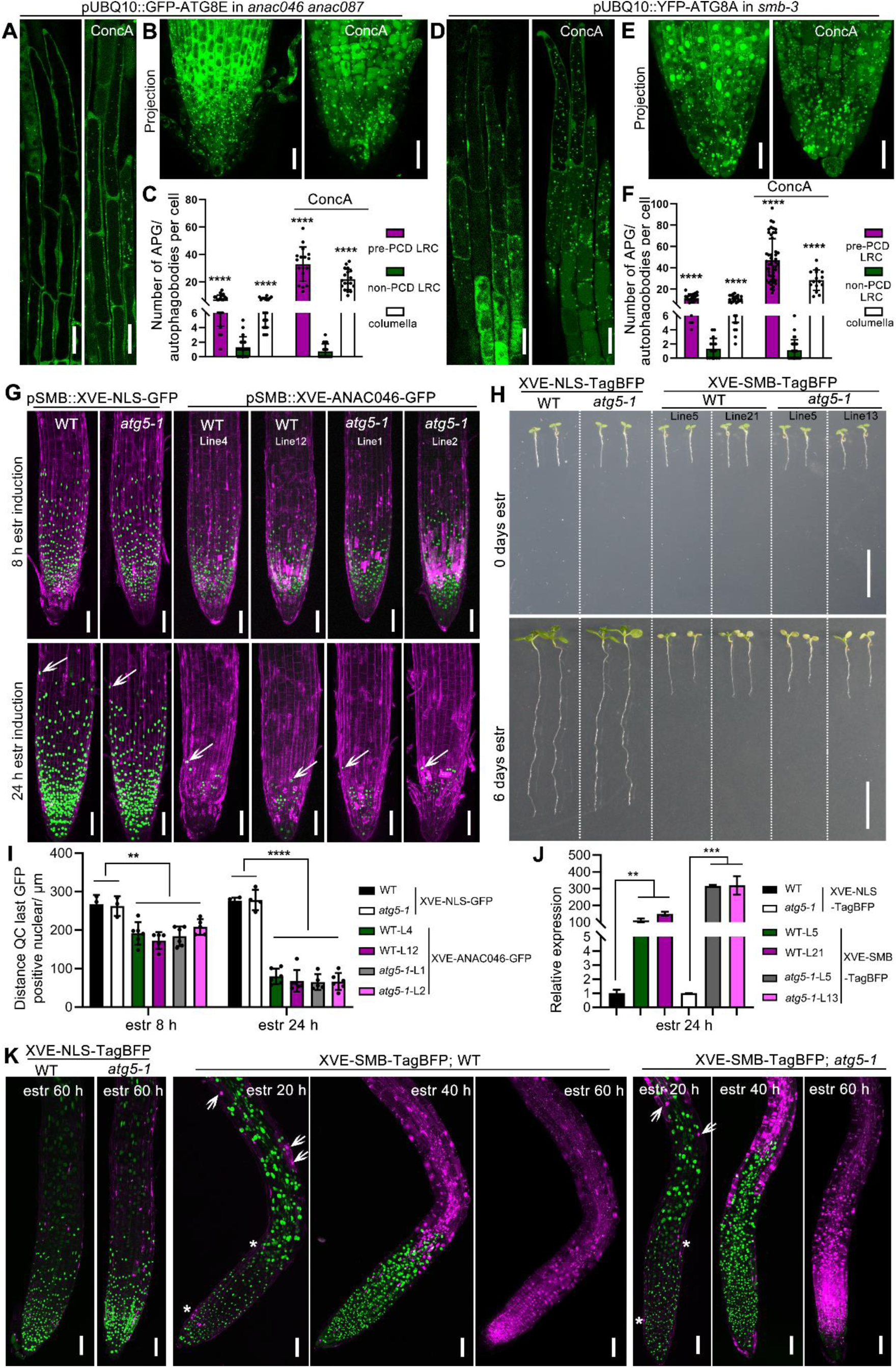
Autophagy occurs independently of established dPCD gene regulatory networks (**A-C**), CLSM of LRC cells from 5 DAG seedlings expressing pUBQ10::GFP-ATG8E in the *anac046 anac087* mutant, longitudinal section of LRC cells (A, left panel), treated with 1 μM ConcA for 8 h (A, right panel), projection of root tip (B, left panel), treated with ConcA (B, right panel). Scale bars are 20 μm. C, Quantification of autophagosomes or autophagic bodies in LRC and columella cells shown in A-B. Results are means ± SD (> 20 counted cells from several separate seedlings). **** indicates a significant difference (*t* test, P < 0.0001). (**D-F**), CLSM of LRC cells from 5 DAG seedlings expressing pUBQ10::YFP-ATG8A in the *smb-3* mutant, longitudinal section of LRC cells (D, left panel), treated with 1 μM ConcA for 8 h (D, right panel), projection of root tip (E, left panel), treated with ConcA (E, right panel). Scale bars are 20 μm. F, Quantification of autophagosomes or autophagic bodies in LRC and columella cells shown in D-E. Results are means ± SD (> 20 counted cells from several separate seedlings). **** indicates a significant difference (*t* test, P < 0.0001). (**G**), CLSM of root tips from 5 DAG seedlings pSMB::XVE-NLS-GFP in wild type, in *atg5-1*, and pSMB::XVE-ANAC046-GFP in wild type and *atg5-1* mutant. Two independent lines are shown, stained with PI (magenta). 8 h after estradiol induction (z-stack projection, up row), 24 h after estradiol induction (z-stack projection, down row). White arrows indicate the position of the last living root cap cells. Scale bars are 50 μm. (**H**), Macroscopic appearance of estradiol-induction of SMB-TagBFP driven by promoter of pH3.3 in wild type and *atg5-1* mutant, estradiol-induction of NLS-TagBFP in wild type or *atg5-1* mutant as control. Two independent lines are shown at 0 day (upper row) and 6 days (bottom row) after transfer to estradiol-containing medium. Overexpression of SMB resulted in root growth arrest in *atg5-1* mutants similar to in the wild type, while the overexpression of NLS-TagBFP did not result in root growth arrest both in *atg5-1* and wild type after 6 days of estradiol induction. Scale bars are 1 cm. (**I**), Quantification of the root cap length of two independent lines each of XVE-ANA046-GFP overexpressors and XVE-NLS-GFP overexpressors shown in G. At least 5 roots of each line were quantified. Results shown are means ± SD. Each line of XVE-ANAC046-GFP is significantly different from line of XVE-NLS-GFP, both in wild type and *atg5-1* mutant, but there is no difference between wild type and *atg5-1*. (*t* test; ** p<0.01, **** p<0.0001). (**J**), As indicated by RT-qPCR analysis, a 24-hour induction by estradiol caused strong expression of SMB in two independent lines of XVE-TagBFP in wild type and *atg5-1* mutant, XVE-NLS-TagBFP in wild type and *atg5-1* as control. Results shown are means ± SD (*t* test; ** p<0.01, *** p<0.001). (**K**), CLSM of root tips from 5 DAG seedlings of pH3.3::XVE-NLS-TagBFP and pH3.3::XVE-SMB-TagBFP in wild type and *atg5-1* mutant at different timepoints after estradiol induction, stained with the PI (magenta). 16 h after estradiol induction, ectopic cell death was detected indicated by white arrows. PCD of root cap cells is indicated by white asterisks. 40 h after estradiol induction abundand cell death occurred above the root meristem. 60 h after estradiol induction the whole root tip is dead and stained with PI. The signal of TagBFP is shown in green. Scale bars are 50 μm.

Conversely, we tested whether the ectopic PCD phenotype caused by inducible misexpression of ANAC046^10^ depends on a functional autophagy pathway. We transformed an estradiol-inducible XVE-ANAC046-GFP expression cassette controlled by the pSMB promoter into *atg5-1* mutants and wild type. Two independent lines in each ecotype were selected based on the GFP signal upon estradiol treatment. 8 h after estradiol induction, we observed both a strong GFP signal in the root cap, as well as an increased number of PI-stained root cap cells in the wild type and the *atg5-1* mutant. This indicates that ANAC046 misexpression under the pSMB promoter is sufficient to cause a precocious wide-spread dPCD in the root cap (Figure 5G, upper row, and I). 24 h after estradiol treatment this phenotype was exacerbated (Figure 5G, down row, and I). Importantly, we did not observe any difference in the effect of ANAC046 expression in the wild type and in *atg5-1* mutants, suggesting that ANAC046-induced cell death in the root cap does not depend on a functional autophagy pathway.

Inducible misexpression of SMB leads to ectopic cell death in the entire plant^10^, and the *smb-3* mutant shows a delayed and aberrant root cap cell death devoid of cell corpse clearance^6^. To investigate if the increased autophagic flux in PCD-preparing root cap cells depends on SMB, we introgressed the YFP-ATG8A marker into the *smb-3* mutant. However, the increased autophagic activity in mature LRC cells in comparison with “non-PCD” LRC cells still occurred in *smb-3* mutants (Figure 5D-F), suggesting that increased autophagic flux prior to root cap PCD is not regulated by SMB. Conversely, to test if the ectopic cell death caused by SMB misexpression depends on the autophagy pathway, we transformed an estradiol-inducible SMB-TagBFP construct controlled by the ubiquitous pH3.3/HTR5 promoter^33^ into the wild type and *atg5-1* mutants. Two independent lines in each ecotype were selected based on inducible transcription of SMB (Figure 5J) and analyzed for their ectopic cell death phenotype after induction. As control, we generated and inducible NLS-TagBFP construct under the same promoter and transformed it into the wild type and *atg5-1* mutants. In the wild type background, ectopic cell death observed in SMB misexpression seedlings coincided with root growth arrest and seedling death upon estradiol treatment (Figure 5H). Microscopic analysis revealed first cells succumbing to ectopic cell death at 20 h after estradiol application. After 40 h cell death had spread throughout the root elongation zone, and at 60 h the entire root tip had died (Figure 5K). The NLS-TagBFP lines showed neither root growth arrest nor cell death upon estradiol treatment (Figure 5K). In the *atg5-1* mutant background, we did not observe any difference in root growth arrest or ectopic cell death when compared to the wild type (Figure 5H-K).

In sum, increased autophagic flux in root cap cells preparing for dPCD occurs independently of the key PCD regulators SMB, ANAC046 and ANAC087, and the ectopic cell death caused by misexpression of these transcription factors does not depend on functional autophagy. These findings suggest that autophagy controls PCD in proximal root cap cells via parallel pathway that operates independently of the established PCD-promoting NAC transcription factors.

## Discussion

Our investigations of autophagy reporters demonstrate an increase in autophagic activities in mature root cap cells prior to the execution of dPCD. This increase can be observed in both the distal LRC (close to the elongation zone) as well as the proximal root cap (columella and adjacent proximal LRC cells). Analysis of several *atg* mutants revealed an unexpectedly differentiated role of autophagy in root cap PCD: Despite the disruption autophagic flux prior to PCD in the distal LRC, PCD occurred timely and in a fashion indistinguishable of from the wild type. However, in the proximal root cap of *atg* mutants, PCD execution was significantly delayed, leading to an increased number of surviving root cap cells during and after their shedding into the rhizosphere. The long-lived autophagy-deficient proximal root cap cells displayed a unique vacuolar morphology and when they finally did die, there was no sign of post-mortem corpse clearance that degrades cytoplasmic contents in wild-type root cap cells. Our investigations did not reveal any interaction between autophagy and known root cap dPCD gene regulatory networks controlled by the NAC transcription factors SMB, ANC046, and ANAC087, suggesting that autophagy acts in an independent parallel pathway.

Our finding that the execution of distal LRC PCD is not dependent on autophagy is in line with the observation that autophagy-defective mutants in Arabidopsis do not show detrimental defects in other developmental processes involving PCD, such as xylem tracheary element differentiation, seed development, or tapetum degeneration^13^. Recently, a rice (*Oryza sativa*) *atg7* mutant has been reported to show defects in pollen maturation and anther dehiscence resulting in reduced male fertility^34^. The rice phenotype might be caused by an autophagy-dependent differentiation and metabolic reprogramming in the anther tapetum, which would imply divergent species-specific roles of autophagy in this tissue.

Autophagy has been implicated in the promotion of xylem PCD^35^. However, not only PCD execution, but also earlier steps of tracheary element differentiation such as formation of a large central vacuole and secondary cell wall formation were compromised in autophagy-deficient cell cultures. This would imply that autophagy promotes xylem differentiation upstream of dPCD, rather than cell death execution itself. Similarly, in suspensor cells of *in-vitro* cultivated Norway spruce (*Piecea abies*) embryos, suppression of autophagy caused cellular differentiation defects including a failure to generate large central vacuoles and the typical anisotropic cell expansion. These differentiation defects were associated with a switch from the regular, autolytic suspensor PCD to a rapid, necrosis-like cell death mode devoid of cell corpse clearance^24^. We show that Arabidopsis ATG-deficient proximal root cap cells also exhibited an altered vacuolar morphology and an inhibition of post-mortem cell corpse autolysis. However, *atg* mutant root cap cells showed a substantial extension of proximal root cap life span instead of a switch to a necrotic-like rapid cell death mode. Furthermore, root cap differentiation does not depend on functional autophagy, as indicated by the wild type-like morphogenesis and differentiation-induced statolith formation^30,31^ in ATG-deficient root cap cells. Conversely, autophagy increase in mature root cap cells is not part of the SMB-dependent dPCD preparation program: In the *smb* mutant, which is delayed in root cap differentiation^7^ and unable to undergo a regular root cap dPCD^6^, autophagic flux occurs as in the wild type. These results suggest that autophagy and cellular differentiation are regulated by independent pathways in the Arabidopsis root cap.

Interestingly, statolith degradation in columella cells prior to PCD was not affected in 5-day old *atg* mutants, although statoliths appeared smaller and more dispersed. Seeds of the rice *Osatg7-1* mutant were found to be smaller and lower in starch content due to an abnormal activation of starch degradation pathways in the endosperm during seed maturation^36^. Possibly, a similar pathway operates in the autophagy-deficient columella. Unlike post-mortem corpse clearance that is severely affected in ATG-deficient columella cells, starch degradation prior to PCD occurs as in the wild type, showing that this process occurs independently of autophagic activities.

Recently, autophagy has been shown to promote cell death processes associated with carbon starvation in tobacco BY-2 suspension culture cells, in which RNAi silencing of ATG4 reduced cell death rates^37^. Conceivably, carbon starvation is occurring and promoting PCD in detached root cap cells, but not in distal LRC cells, which would be in line with the different roles of autophagy we observed in these two cell types.

In Arabidopsis cell suspension cultures stimulated to differentiate into xylem cell types, depletion of METACASPASE9 caused increased levels of autophagy in cells that undergo tracheary element (TE) differentiation, but did not affect PCD in these cells. However, an increased cell death rate in surrounding non-TE cells that normally stay alive was reported. Downregulation of autophagy by ATG2 depletion specifically in TE-like cells restored viability of non-TE cells, suggesting that autophagy contributes to confine cell death to TE cells^38^. Similarly, upon TE-specific overexpression of ATG5 during ectopically induced xylem-differentiation in Arabidopsis cotyledons, TE differentiation and PCD were not affected, but increased cell death in neighboring non-TE cells was observed^39^. These findings indicate a non-cell autonomous role of autophagy and demonstrate the need to manipulate autophagy using cell-type specific approaches.

Already previously, cell-type specific approaches had been called for to address the specific roles of autophagy in different cell types^40^. In line with this, a recent study found different autophagy activity in root hair and non-root hair cell files of the Arabidopsis root epidermis, indicating differences in autophagic pathway regulation these cell types (doi: https://doi.org/10.1101/2021.12.07.471480). Our CRISPR-TSKO root-cap specific autophagy knock-out approach clearly shows that the PCD-promoting role of autophagy is controlled autonomously within the root cap. Such approaches will be a powerful addition to the autophagy toolbox, and able to address the roles of autophagy in specific cell types, tissues, or organs while avoiding pleiotropic or secondary effects of conventional knockouts or pharmacological treatments.

## Methods

### Plant materials and growth conditions

The *Arabidopsis thaliana atg2-2* mutant allele (EMS, Gln803stop) was reported by^18^; *atg5-1* (SAIL_129_B07) was reported by^17^; *atg7-2* (GABI_655B06) was reported by^41^, *smb-3* was reported by^6^, *anac046 anac087* was reported by^10^. The Arabidopsis lines pUBQ10::YFP-ATG8A, p35S::mCherry-ATG8E^26^ and pSMB::NLS-GFP^6^ were introduced into *atg5-1* and/or *atg2-2*, or *smb-3* mutant by crossing, respectively. The line p35S::YFP-mCherry-NBR1 was reported by^29^. The pUBQ10::GFP-ATG8E was introduced into *anac046 anac087* by dipping. pUBQ10::ToIM was reported previously^10^.

All Arabidopsis seedlings were grown vertically on 1/2 Murashige and Skoog (MS, Duchefa Biochemie) medium (0.1 g/L MES, pH 5.8 [KOH], and 0.8% plant agar) in a continuous light at 21°C before analysis, except where noted.

### Cloning

Golden Gate entry modules pGG-A-pSMB-B, pGG-B-Linker-C, pGG-C-Cas9-D, pGG-D-P2A-GFP-NLS-E, pGG-E-G7T-F, pGG-F-pATU6-26-AarI-G, were reported previously^32^. These entry modules were assembled in pFASTR-AG, resulting in the destination vector pFASTR-pSMB-Cas9-P2A-GFP-NLS-pATU6-26-AarI. Fragment gRNA1-pATU6-26-gRNA2 (ATG2 target) and gRNA1-pATU6-26-gRNA2 (ATG5 target) were amplified by PCR using the following primers, p43/p44 for ATG2, p41/42 for ATG5, and These purified PCR fragments were inserted into pFASTR-pSMB-Cas9-P2A-GFP-NLS-pATU6-26-AarI destination vector via a Golden Gate reaction. The resulting vectors were named pSMB-Cas9;ATG2 and pSMB-Cas9;ATG5.

Golden Gate entry modules pGG-A-pUBQ10-B, pGG-B-mGFP-C, pGG-C-ATG8E-D, pGG-D-Linker-E, pGG-E-tHSP18.2M-F, pGG-F-linkerII-G were collected from PSB plasmids stock. And these entry modules were assembled in pFASTRK-AG, resulting in the expression vector pFASTRK-pUBQ10::GFP-ATG8E.

Gateway entry modules L4-pSMB-R1, L1-ANAC046-L2, and R2-GFP-L3 were described previously^6,10^. L4-pSMB-XVE-R1, L4-pH3.3/HTR5 -XVE-R1, L1-NLS-GFP-L2, L1-NLS-TagBFP-L2, L1-SMB-L2 and R2-TagBFP-L3 were ordered from the PSB plasmid stock (https://gatewayvectors.vib.be). The ATG5 coding sequence was amplified by PCR with primers p214 and p215, using 4-day old seedling cDNA as template and inserted into pDONR221 via BP reaction, resulting in entry vector, L1-ATG5-L2. Entry vectors L4-pSMB-XVE-R1, L1-ANAC046-L2 and R2-GFP-L3 were assembled into pFASTRK-34GW destination vector via LR reaction. The result vector was pFASTRK-pSMB::XVE-ANAC046-GFP. L4-pSMB-XVE-R1, L1-NLS-GFP-L2 were assembled into pFASTRK-24GW destination vector via LR reaction. The resulting vector was pFASTRK-pSMB::XVE-NLS-GFP. L4-pH3.3/HTR5 -XVE-R1, L1-SMB-L2 and R2-TagBFP-L3 were assembled into pFASTGB-34GW destination vector via LR reaction. The resulting vector was pFASTGB-pH3.3/HTR5::XVE-SMB-TagBFP. L4-pH3.3/HTR5 -XVE-R1, L1-NLS-TagBFP-L2 were assembled into pFASTGB-24GW destination vector via LR reaction. The resulting vector was pFASTGB-pH3.3/HTR5::XVE-NLS-TagBFP. L4-pSMB-R1, L1-ATG5-L2 and R2-GFP-L3 were assembled into pFASTGB-34GW destination vector via LR reaction. The resulting vector was pFASTGB-pSMB::ATG5-GFP.

All PCRs for cloning was performed with Phusion high-fidelity DNA polymerase (Thermo Scientific). All entry vectors were sequenced by Eurofins Scientific using the Mix2Seq or TubeSeq services. All primers are listed in Table S1.

### Staining and imaging

Confocal imaging was done on and LSM710 (Zeiss) using the Plan Apochromat 20× objective (numerical aperture 0.8), or an SP8X (Leica) using a 40x (HC PL APO CS2, NA=1.10) water immersion objective, unless stated otherwise.

For the FDA-PI viability staining, seedlings were mounted on a glass slide in FDA solution (1 μL dissolved FDA stock solution [2 mg in 1 ml acetone] in 1 ml 0.5 1/2 MS) supplemented with 10 μg/ml PI. FDA and PI were imaged simultaneously on the LSM710 (Zeiss) using the 488-nm laser line to excite FDA and the 561-nm laser line to excite PI. Emission was detected between 500 and 550 nm and between 600 and 700 nm for FDA and PI, respectively. For the FDA-FM4-64 staining, seedlings were mounted on a glass slide in FDA solution (1 μL dissolved FDA [2 mg in 1 ml acetone] in 1 ml 1/2 MS) supplemented with 4 μg/ml FM4-64. FM4-64 was imaged on the LSM710 (Zeiss) using the 561-nm laser line to excite FM4-64. Emission was detected between 600 and 700 nm for FM4-64.

In the inducible overexpression lines, seedlings were sprayed with 50 μg/ml estradiol or DMSO (mock) and then mounted on a glass slide in 1/2 MS supplemented with 10 μg/ml PI at different timepoints. GFP/ TagBFP and PI were detected simultaneously in the same track on the LSM710 (Zeiss). GFP was excited by the 488-nm line of the argon laser and detected between 500 and 550 nm in channel 1, or TagBFP was excited by 405-nm line and detected between 450 and 510 nm in channel 1, whereas PI was excited by the 561-nm line and detected between 590 and 700 nm in channel 2. For the overnight time course, 20 h induced seedlings were transferred to a Lab-Tek chamber and covered with an agar slab (1/2 MS) supplemented with 10 μg/ml estradiol and 10 μg/ml PI. Roots were imaged at different timepoints.

For the reporter lines of pUBQ10::YFP-ATG8A, pUBQ10::GFP-ATG8E, YFP-mCherry-NBR1, seedlings were mounted on a glass slide in 1/2 MS medium and imaged on SP8 (Leica). YFP was excited by the 514-nm line of the argon laser and detected between 525 and 580 nm. GFP was excited by the 488-nm line of the argon laser and detected between 500 and 550 nm. The signal of mCherry was excited by the 561-nm line and detected between 600 nm and 700 nm. Imaging of ToIM was performed as described before^10^.

For starch staining, 5 days old seedlings were stained with Lugol’s iodine solution (Sigma-Aldrich, 62650) for 5 min, washed in distilled water for 1 min and then mounted with clearing solution (chloral hydrate: glycerol: water=8:1:3). Samples were imaged on an Olympus BX51 microscope with a 40× DIC objective.

### Transmission Electron Microcopy (TEM)

Root tips of 5 days old and 14 days old seedlings of Arabidopsis thaliana Col-0 and atg2-2 were excised, immersed in 20% (w/v) BSA and frozen immediately in a high-pressure freezer (Leica EM ICE; Leica Microsystems, Vienna, Austria). Freeze substitution was carried out using a Leica EM AFS (Leica Microsystems) in dry acetone containing 1% (w/v) OsO4 and 0.2% glutaraldehyde over a 4-days period as follows: - 90°C for 54 hours, 2°C per hour increase for 15 hours, -60°C for 8 hours, 2°C per hour increase for 15 hours, and -30°C for 8 hours. Samples were then slowly warmed up to 4°C, infiltrated stepwise over 3 days at 4°C in Spurr’s resin and embedded in capsules. The polymerization was performed at 70 °C for 16 h. Ultrathin sections were made using an ultra-microtome (Leica EM UC6) and post-stained in in a Leica EM AC20 for 40 min in uranyl acetate at 20 °C and for 10 min in lead stain at 20 °C. Sections were collected on formvar-coated copper slot grids.

Grids were viewed with a JEM1400plus transmission electron microscope (JEOL, Tokyo, Japan) operating at 80 kV.

## Abbreviations

PCD: programmed cell death
dPCD: developmental PCD
LRC: lateral root cap
RCD: regulated cell death
ConcA: Concanamycin A
FDA: fluorescein diacetate
PI: propidium iodide
TE: tracheary element
ToIM: tonoplast integrity marker.

## Accession Numbers

Gene models used in this article can be found in the Arabidopsis Genome Initiative database under the following accession numbers: ATG2 (AT3G19190); ATG5 (AT5G17290); ATG7 (AT5G45900); ATG8A (AT4G21980); ATG8E (AT2G45170); SMB (AT1G79580); ANAC046 (AT3G04060); ANAC087 (AT5G18270).

## Acknowledgments

We thank Peter Bozhkov (Department of Chemistry and Biotechnology, Uppsala BioCenter, Swedish University of Agricultural Sciences, Sweden) for the seeds of *atg5-1, atg7-2* and GFP-ATG8A in *atg5-1*; Celine Masclaux-Daubresse for seeds of p35S::mCherry-ATG8E (originally) from Romain Lebars; Matyas Fendrych for the seeds of pUBQ10::YFP-ATG8A in Col-0. This research was financially supported by the European Research Council (ERC) StG PROCELLDEATH 639234 and CoG EXECUT.ER 864952 (MKN), and by a postdoctoral fellowship of the University of Ghent Bijzonder Onderzoeksfonds (BOF) (QF).

## Supplemental Material

### Supplemental Movie legends

**Movie S1**. Numerous YFP-ATG8A-positive autophagic bodies are present in wild type columella cells.

The root tip of a wild type seedling at 5 DAG expressing pUBQ10-YFP-ATG8A treated with ConcA for 8 h. Scale bar is 20 μm.

**Movie S2**. The autophagy-deficient *atg5-1* mutant columella cells contain no autophagic bodies but large YFP-ATG8A protein aggregates.

The root tip an *atg5-1* mutant seedling at 5 DAG expressing pUBQ10-YFP-ATG8A treated with ConcA for 8 h. Scale bar is 20 μm.

### Supplemental Figures S1 – S5

**Figure S1.**
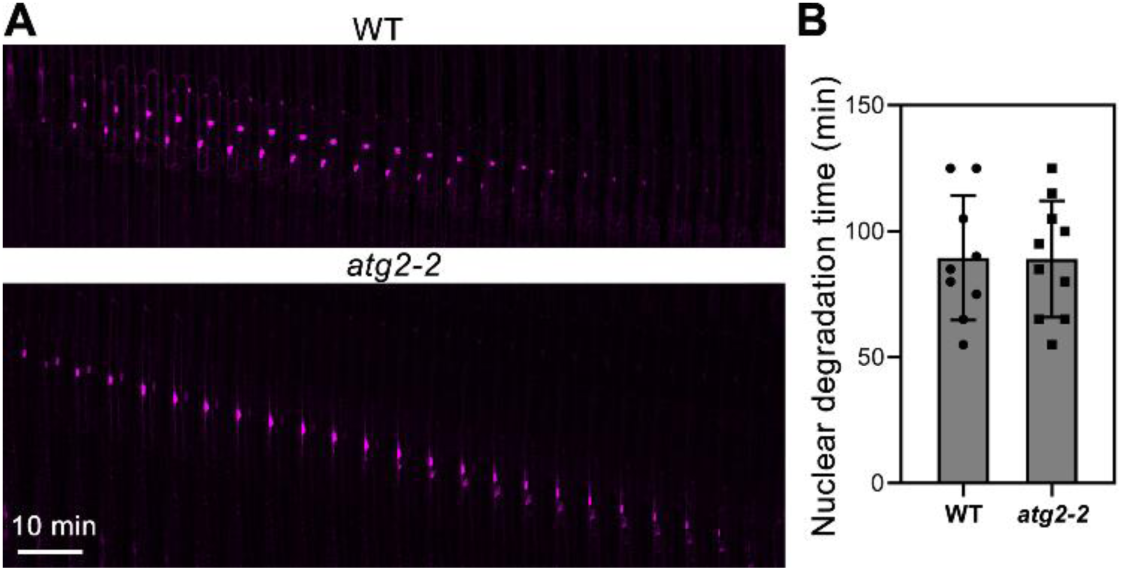
Nuclear degradation in distal LRC cells is indistinguishable in the wild type and the *atg2-2* mutant. (**A**), Kymograph showing nuclear degradation in *atg2-2* mutant compared with the wild type. Cells were imaged in 5 min intervals for 2 h after staining with PI (magenta). (**B**), Quantification of the time of nuclear degradation shown in (A). Results are means ± SD. There is not significant difference in wild type and *atg2-2* mutant (Student’s *t* test, p > 0.05).

**Figure S2.**
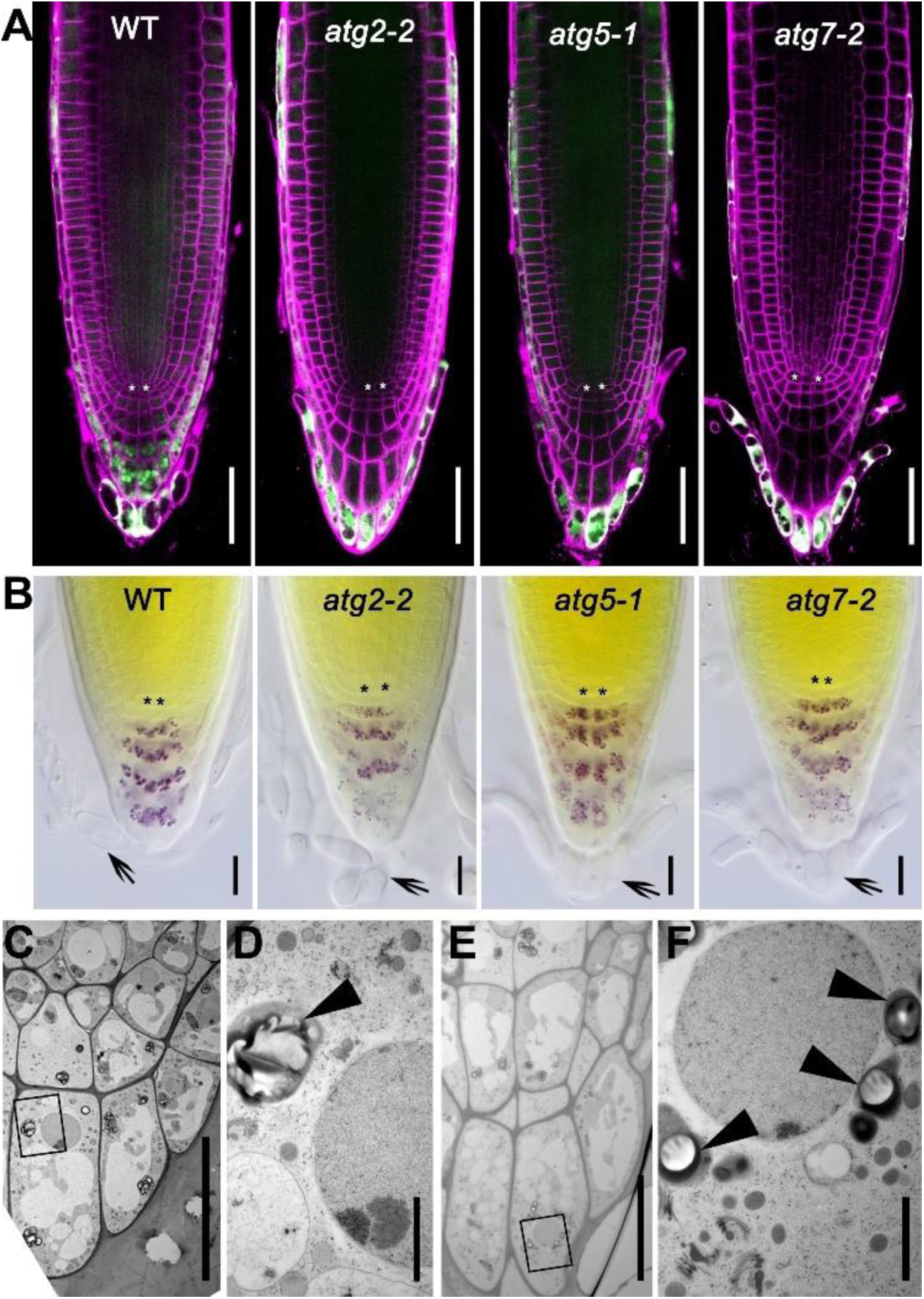
Analysis of root tip morphology and statolith formation in columella cells of the wild type and *atg* mutants shows that root cap development and differentiation occur independent of autophagy. (**A**), Longitudinal sections of *Arabidopsis* roots at 5 DAG pulse labeled with FDA (green) and FM4-64 (magenta). White asterisks indicate quiescent center cells (QC). Scale bars are 50 μm. (**B**), Iodine-stained roots of the wild type and *atg* mutants in 5 DAG seedlings. Black asterisks indicate quiescent center cells (QC). Arrows point to the outermost root cap layer in which statoliths are degraded. Scale bars are 20 μm. (**C**), TEM micrograph of columella cells in a wild-type seedling at 5 DAG. Scale bar is 20 μm. (**D**), Close-up area of area indicated in (C). Black arrowhead indicates statoliths containing bright starch granules. Scale bar is 2 μm. (**E**), TEM micrograph of columella cells in an *atg2-2* mutant seedling at 5 DAG. Scale bar is 20 μm. (**F**), Close-up area of area indicated in (E). Black arrowheads indicate small statholiths with starch granules. Scale bar is 2 μm.

**Figure S3.**
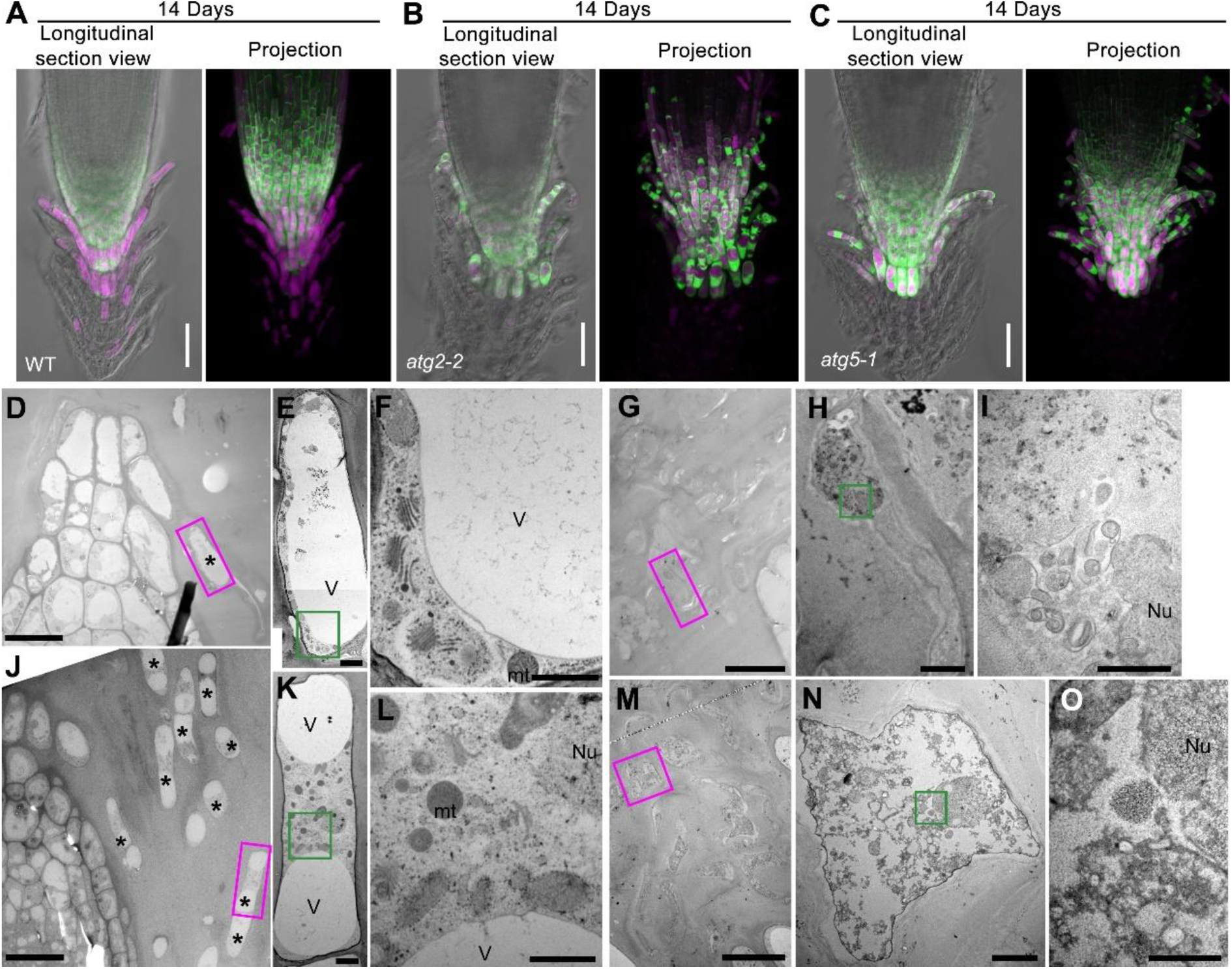
Autophagy regulates PCD onset in proximal LRC cells and corpse clearance in columella cells. (**A-C**), pUBQ10::ToIM expression in the wild type (A), *atg2-2* (B), and *atg5-1* (C) was analyzed at 14 DAG. Cytoplasmic EGFP signal is shown in green, and vacuolar mRFP signal is shown in magenta. Scale bars are 50 μm. (**D-O**), TEM of wild type (D-I) and *atg2-2* (J-O) seedlings at 14 DAG. Closeups of the wild type (E) and *atg2-2* (K) show viable proximal LRC cells that are indicated by magenta frames in (D) and (J), respectively; closeups of the wild type (F) and *atg2-2* (L) indicated by the green frames in (E) and (K), respectively. Closeups of the wild type (H) and *atg2-2* (N) showing dead columella cells indicated by magenta frames in (G) and (M), respectively; closeups of the wild type (I) and *atg2-2* (O) indicated by green frames in (H) and (N), respectively. The asterisks indicate alive proximal LRC cells. V: vacuole, mt: mitochondria, Nu: nucleus or nuclear remnant. Scale bars are 20 μm for D, G, J and M, 2 μm for E, H, K, and N, 1 μm for F and L, and 500 nm for I and O.

**Figure S4.**
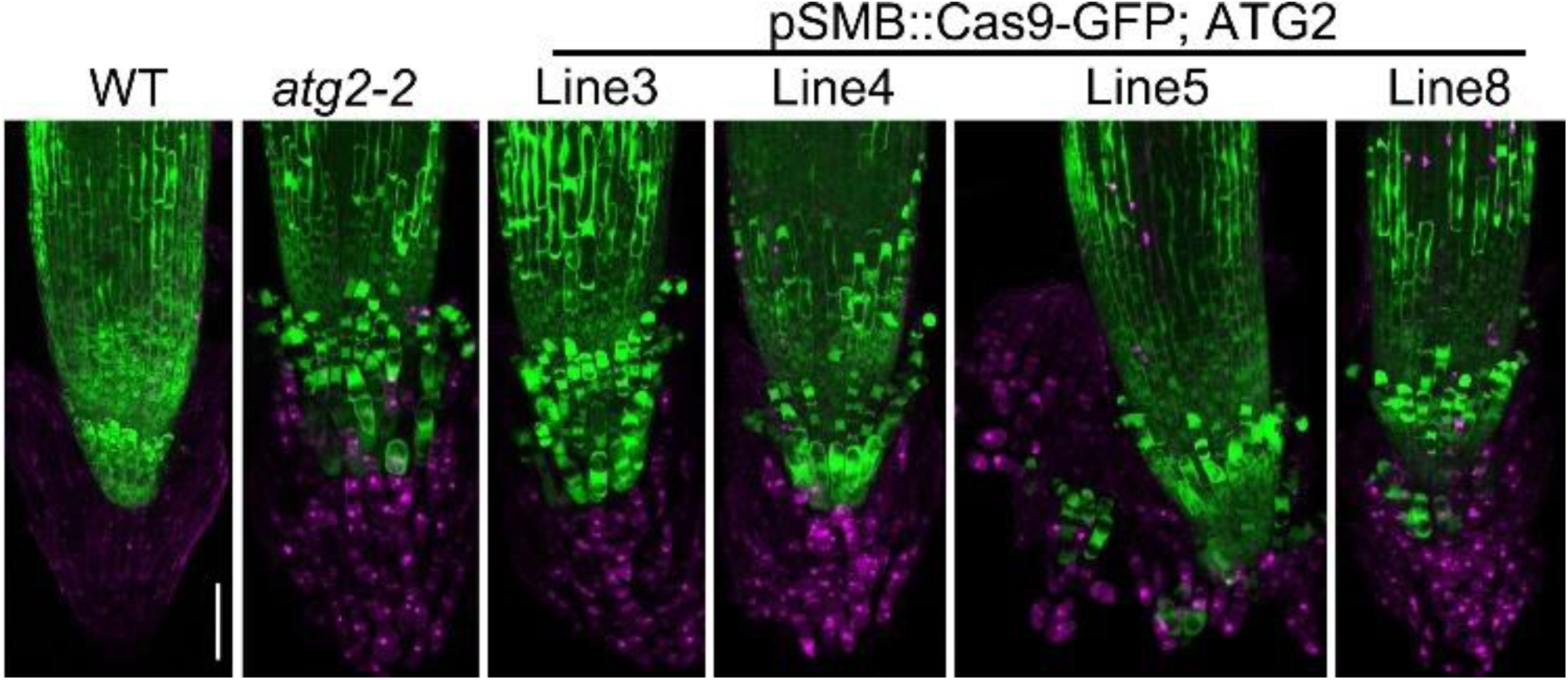
Root cap specific CRISPR-TSKO of ATG2 mimics the *atg2-2* phenotype. CLSM (z-stack projection) of root tips of seedlings carrying a pSMB::NLS-GFP construct at 14 DAG. From left to right: wild type (WT), *atg2-2*, and four independent lines carrying a root cap-specific CRISPR-TSKO construct targeting ATG2 (pSMB::Cas9-GFP;ATG2), pulse labeled with FDA (green) and PI (magenta). Scale bar is s50 μm.

**Figure S5.**
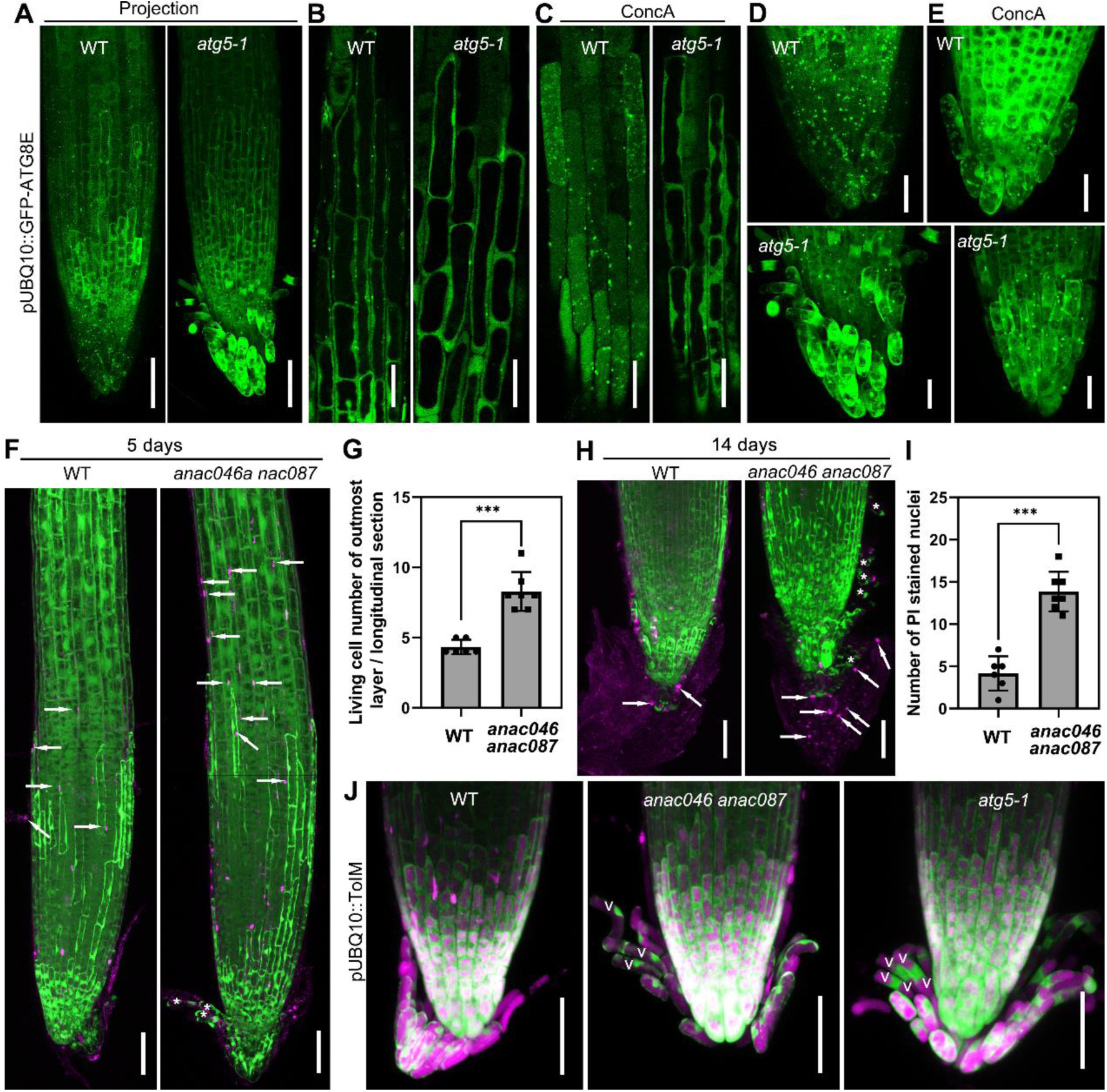
Expression of pUBQ10::GFP-ATG8E in the wild type and *atg5-1*, and phenotypic analyses of the *anac046 anac087* mutant. (**A**), CLSM (z-stack projection) of root tips of wild type and *atg5-1* seedlings at 5 DAG expressing GFP-ATG8E. Scale bars are 50 μm. (**B** and **C**), Distal LRC cells of wild type and *atg5-1* seedlings at 5 DAG expressing GFP-ATG8E (B), treated with 1 μM ConcA for 8 h (C). Scale bars are 20 μm. (**D** and **E**), Proximal LRC and columella cells of wild type (top row) and *atg5-1* (bottom row) seedlings at 5 DAG expressing GFP-ATG8E (D), treated with 1 μM ConcA for 8 h (E). z-stack projection. Scale bars are 50 μm. (**F**), CLSM (z-stack projection) of root tips wild type and *anac046 anac087* mutant seedlings at 5 DAG, stained with FDA and PI. White arrows point to cell corpses in distal LRC cells. Asterisks indicate viable proximal LRC cells. Scale bars are 50 μm. (**G**), Quantification of viable cells in the outermost root cap cell layer of wild type and *anac046 anac087* mutant seedlings at 5 DAG counted on longitudinal root sections. Results are means ± SD. *** indicates a significant difference (*t* test, P < 0.001). (**H**), CLSM (z-stack projection) of root tips from 14 DAG seedlings of wild type and *anac046 anac087* mutant, stained with FDA and PI. White arrows point cell corpse in columella cells. Asterisks point alive proximal LRC cells. Scale bars are 50 μm. (**I**), Quantification of PI-positive (dead but non-degraded) cells in the outermost root cap cell layer of wild type and *anac046 anac087* mutant seedlings, as shown in (H). Results are means ± SD. *** indicates a significant difference (*t* test, P < 0.001). (**J**), Tonoplast integrity marker in wild type, *anac046 anac087* mutant and *atg5-1* mutant seedlings at 5 DAG. V: vacuole. Scale bars are 50 μm.

## References

1. Galluzzi, L., Vitale, I., Aaronson, S.A., Abrams, J.M., Adam, D., Agostinis, P., Alnemri, E.S., Altucci, L., Amelio, I., Andrews, D.W., et al. (2018). Molecular mechanisms of cell death: recommendations of the Nomenclature Committee on Cell Death 2018. Cell death and differentiation 25, 486–541. 10.1038/s41418-017-0012-4.

2. Huysmans, M., Lema, A.S., Coll, N.S., and Nowack, M.K. (2017). Dying two deaths - programmed cell death regulation in development and disease. Current opinion in plant biology 35, 37–44. 10.1016/j.pbi.2016.11.005.

3. Pitsili, E., Phukan, U.J., and Coll, N.S. (2020). Cell Death in Plant Immunity. Cold Spring Harb Perspect Biol 12. 10.1101/cshperspect.a036483.

4. Daneva, A., Gao, Z., Van Durme, M., and Nowack, M.K. (2016). Functions and Regulation of Programmed Cell Death in Plant Development. Annual review of cell and developmental biology 32, 441–468. 10.1146/annurev-cellbio-111315-124915.

5. Rost, T.L. (2011). The organization of roots of dicotyledonous plants and the positions of control points. Annals of botany 107, 1213–1222. 10.1093/aob/mcq229.

6. Fendrych, M., Van Hautegem, T., Van Durme, M., Olvera-Carrillo, Y., Huysmans, M., Karimi, M., Lippens, S., Guerin, C.J., Krebs, M., Schumacher, K., and Nowack, M.K. (2014). Programmed cell death controlled by ANAC033/SOMBRERO determines root cap organ size in Arabidopsis. Current biology : CB 24, 931–940. 10.1016/j.cub.2014.03.025.

7. Bennett, T., van den Toorn, A., Sanchez-Perez, G.F., Campilho, A., Willemsen, V., Snel, B., and Scheres, B. (2010). SOMBRERO, BEARSKIN1, and BEARSKIN2 regulate root cap maturation in Arabidopsis. The Plant cell 22, 640–654. 10.1105/tpc.109.072272.

8. Kumpf, R.P., and Nowack, M.K. (2015). The root cap: a short story of life and death. Journal of experimental botany 66, 5651–5662. 10.1093/jxb/erv295.

9. Shi, C.L., von Wangenheim, D., Herrmann, U., Wildhagen, M., Kulik, I., Kopf, A., Ishida, T., Olsson, V., Anker, M.K., Albert, M., et al. (2018). The dynamics of root cap sloughing in Arabidopsis is regulated by peptide signalling. Nature plants 4, 596–604. 10.1038/s41477-018-0212-z.

10. Huysmans, M., Andrade Buono, R., Skorzinski, N., Cubria Radio, M., De Winter, F., Parizot, B., Mertens, J., Karimi, M., Fendrych, M., and Nowack, M.K. (2018). ANAC087 and ANAC046 control distinct aspects of programmed cell death in the Arabidopsis columella and lateral root cap. The Plant cell. 10.1105/tpc.18.00293.

11. Olvera-Carrillo, Y. M.V.B., Tom Van Hautegem, Matyáš Fendrych3,, M.H., Maria Simaskova, M.v.D., Pierre Buscaill, Susana Rivas, Nuria S. Coll, Frederik Coppens,, and Steven Maere, a.M.K.N. (2015). A Conserved Core of Programmed Cell Death Indicator Genes Discriminates Developmentally and Environmentally Induced Programmed Cell Death in Plants1[OPEN]. Plant physiology 169, 2684–2699.

12. Zhang, H., and Baehrecke, E.H. (2015). Eaten alive: novel insights into autophagy from multicellular model systems. Trends Cell Biol 25, 376–387. 10.1016/j.tcb.2015.03.001.

13. Marshall, R.S., and Vierstra, R.D. (2018). Autophagy: The Master of Bulk and Selective Recycling. Annual review of plant biology 69, 173–208. 10.1146/annurev-arplant-042817-040606.

14. Li, F., and Vierstra, R.D. (2012). Autophagy: a multifaceted intracellular system for bulk and selective recycling. Trends in plant science 17, 526–537. 10.1016/j.tplants.2012.05.006.

15. Liu, Y., and Bassham, D.C. (2012). Autophagy: pathways for self-eating in plant cells. Annual review of plant biology 63, 215–237. 10.1146/annurev-arplant-042811-105441.

16. Doelling, J.H., Walker, J.M., Friedman, E.M., Thompson, A.R., and Vierstra, R.D. (2002). The APG8/12-activating enzyme APG7 is required for proper nutrient recycling and senescence in Arabidopsis thaliana. The Journal of biological chemistry 277, 33105–33114. 10.1074/jbc.M204630200.

17. Thompson, A.R., Doelling, J.H., Suttangkakul, A., and Vierstra, R.D. (2005). Autophagic nutrient recycling in Arabidopsis directed by the ATG8 and ATG12 conjugation pathways. Plant physiology 138, 2097–2110. 10.1104/pp.105.060673.

18. Wang, Y., Nishimura, M.T., Zhao, T., and Tang, D. (2011). ATG2, an autophagy-related protein, negatively affects powdery mildew resistance and mildew-induced cell death in Arabidopsis. The Plant journal : for cell and molecular biology 68, 74–87. 10.1111/j.1365-313X.2011.04669.x.

19. Ustun, S., Hafren, A., and Hofius, D. (2017). Autophagy as a mediator of life and death in plants. Current opinion in plant biology 40, 122–130. 10.1016/j.pbi.2017.08.011.

20. Dauphinee, A.N., Denbigh, G.L., Rollini, A., Fraser, M., Lacroix, C.R., and Gunawardena, A. (2019). The Function of Autophagy in Lace Plant Programmed Cell Death. Frontiers in plant science 10, 1198. 10.3389/fpls.2019.01198.

21. Liu, Y., Schiff, M., Czymmek, K., Talloczy, Z., Levine, B., and Dinesh-Kumar, S.P. (2005). Autophagy regulates programmed cell death during the plant innate immune response. Cell 121, 567–577. 10.1016/j.cell.2005.03.007.

22. Yoshimoto, K., Jikumaru, Y., Kamiya, Y., Kusano, M., Consonni, C., Panstruga, R., Ohsumi, Y., and Shirasu, K. (2009). Autophagy negatively regulates cell death by controlling NPR1-dependent salicylic acid signaling during senescence and the innate immune response in Arabidopsis. The Plant cell 21, 2914–2927. 10.1105/tpc.109.068635.

23. Kwon, S.I., Cho, H.J., and Park, O.K. (2010). Role of Arabidopsis RabG3b and autophagy in tracheary element differentiation. Autophagy 6, 1187–1189. 10.4161/auto.6.8.13429.

24. Minina, E.A., Filonova, L.H., Fukada, K., Savenkov, E.I., Gogvadze, V., Clapham, D., Sanchez-Vera, V., Suarez, M.F., Zhivotovsky, B., Daniel, G., et al. (2013). Autophagy and metacaspase determine the mode of cell death in plants. J Cell Biol 203, 917–927. 10.1083/jcb.201307082.

25. Hofius, D., Schultz-Larsen, T., Joensen, J., Tsitsigiannis, D.I., Petersen, N.H., Mattsson, O., Jorgensen, L.B., Jones, J.D., Mundy, J., and Petersen, M. (2009). Autophagic components contribute to hypersensitive cell death in Arabidopsis. Cell 137, 773–783. 10.1016/j.cell.2009.02.036.

26. Le Bars, R., Marion, J., Le Borgne, R., Satiat-Jeunemaitre, B., and Bianchi, M.W. (2014). ATG5 defines a phagophore domain connected to the endoplasmic reticulum during autophagosome formation in plants. Nature communications 5, 4121. 10.1038/ncomms5121.

27. Li, F., Chung, T., Pennington, J.G., Federico, M.L., Kaeppler, H.F., Kaeppler, S.M., Otegui, M.S., and Vierstra, R.D. (2015). Autophagic recycling plays a central role in maize nitrogen remobilization. The Plant cell 27, 1389–1408. 10.1105/tpc.15.00158.

28. Tamura, K., Shimada, T., Ono, E., Tanaka, Y., Nagatani, A., Higashi, S.I., Watanabe, M., Nishimura, M., and Hara-Nishimura, I. (2003). Why green fluorescent fusion proteins have not been observed in the vacuoles of higher plants. The Plant journal : for cell and molecular biology 35, 545–555. 10.1046/j.1365-313x.2003.01822.x.

29. Svenning, S., Lamark, T., Krause, K., and Johansen, T. (2014). Plant NBR1 is a selective autophagy substrate and a functional hybrid of the mammalian autophagic adapters NBR1 and p62/SQSTM1. Autophagy 7, 993–1010. 10.4161/auto.7.9.16389.

30. Hong, J.H., Chu, H., Zhang, C., Ghosh, D., Gong, X., and Xu, J. (2015). A quantitative analysis of stem cell homeostasis in the Arabidopsis columella root cap. Frontiers in plant science 6, 206. 10.3389/fpls.2015.00206.

31. van den Berg, C., Willemsen, V., Hendriks, G., Weisbeek, P., and Scheres, B. (1997). Short-range control of cell differentiation in the Arabidopsis root meristem. Nature 390, 287–289. 10.1038/36856.

32. Decaestecker, W., Buono, R.A., Pfeiffer, M.L., Vangheluwe, N., Jourquin, J., Karimi, M., Van Isterdael, G., Beeckman, T., Nowack, M.K., and Jacobs, T.B. (2019). CRISPR-TSKO: A Technique for Efficient Mutagenesis in Specific Cell Types, Tissues, or Organs in Arabidopsis. The Plant cell 31, 2868–2887. 10.1105/tpc.19.00454.

33. Ingouff, M., Selles, B., Michaud, C., Vu, T.M., Berger, F., Schorn, A.J., Autran, D., Van Durme, M., Nowack, M.K., Martienssen, R.A., and Grimanelli, D. (2017). Live-cell analysis of DNA methylation during sexual reproduction in Arabidopsis reveals context and sex-specific dynamics controlled by noncanonical RdDM. Genes Dev 31, 72–83. 10.1101/gad.289397.116.

34. Hanamata, S., Sawada, J., Ono, S., Ogawa, K., Fukunaga, T., Nonomura, K.I., Kimura, S., Kurusu, T., and Kuchitsu, K. (2020). Impact of Autophagy on Gene Expression and Tapetal Programmed Cell Death During Pollen Development in Rice. Frontiers in plant science 11, 172. 10.3389/fpls.2020.00172.

35. Kwon, S.I., Cho, H.J., Jung, J.H., Yoshimoto, K., Shirasu, K., and Park, O.K. (2010). The Rab GTPase RabG3b functions in autophagy and contributes to tracheary element differentiation in Arabidopsis. The Plant journal : for cell and molecular biology 64, 151–164. 10.1111/j.1365-313X.2010.04315.x.

36. Sera, Y., Hanamata, S., Sakamoto, S., Ono, S., Kaneko, K., Mitsui, Y., Koyano, T., Fujita, N., Sasou, A., Masumura, T., et al. (2019). Essential roles of autophagy in metabolic regulation in endosperm development during rice seed maturation. Sci Rep 9, 18544. 10.1038/s41598-019-54361-1.

37. Teper-Bamnolker, P., Danieli, R., Peled-Zehavi, H., Belausov, E., Abu-Abied, M., Avin-Wittenberg, T., Sadot, E., and Eshel, D. (2021). Vacuolar processing enzyme translocates to the vacuole through the autophagy pathway to induce programmed cell death. Autophagy 17, 3109–3123. 10.1080/15548627.2020.1856492.

38. Escamez, S., Andre, D., Zhang, B., Bollhoner, B., Pesquet, E., and Tuominen, H. (2016). METACASPASE9 modulates autophagy to confine cell death to the target cells during Arabidopsis vascular xylem differentiation. Biol Open 5, 122–129. 10.1242/bio.015529.

39. Escamez, S., Stael, S., Vainonen, J.P., Willems, P., Jin, H., Kimura, S., Van Breusegem, F., Gevaert, K., Wrzaczek, M., and Tuominen, H. (2019). Extracellular peptide Kratos restricts cell death during vascular development and stress in Arabidopsis. Journal of experimental botany 70, 2199–2210. 10.1093/jxb/erz021.

40. Wada, S., Ishida, H., Izumi, M., Yoshimoto, K., Ohsumi, Y., Mae, T., and Makino, A. (2009). Autophagy plays a role in chloroplast degradation during senescence in individually darkened leaves. Plant physiology 149, 885–893. 10.1104/pp.108.130013.

41. Lai, Z., Wang, F., Zheng, Z., Fan, B., and Chen, Z. (2011). A critical role of autophagy in plant resistance to necrotrophic fungal pathogens. The Plant journal : for cell and molecular biology 66, 953–968. 10.1111/j.1365-313X.2011.04553.x.

